# Near-neutral pH increased *n*-caprylate production in a microbiome with product inhibition of methanogenesis

**DOI:** 10.1101/2022.04.01.486710

**Authors:** Rodolfo Palomo-Briones, Jiajie Xu, Catherine M. Spirito, Joseph G. Usack, Lauren H. Trondsen, Juan J. L. Guzman, Largus T. Angenent

**Author notes:** Corresponding author. Schnarrenbergstr. 94-96, 72076 Tübingen, Germany. These authors contributed equally (Rodolfo Palomo-Briones, Jiajie Xu).

## Abstract

The pH is a critical parameter in chain-elongating bioreactors, affecting: **(1)** the concentration of inhibitory undissociated carboxylic acids, which in turn affects the efficiency of product extraction; **(2)** the thermodynamics; and **(3)** the kinetics. Here, we examined the effect of five different pH levels (5.5 to 7.0) on *n*-caprylate (C8) production using an anaerobic sequencing batch reactor (ASBR) with continuous membrane-based liquid-liquid extraction (pertraction). We found that the product spectrum was directed by pH: mildly acidic pH (5-6) led to *n*-caproate (C6) production, while near-neutral and neutral pH (6.75-7) favored *n*-caprylate production. In particular, the pH of 6.75 led to the maximum values of volumetric *n*-caprylate production rate (75.6 ± 0.6 mmol C L^−1^ d^−1^; 0.06 g L^−1^ d^−1^) and *n*-caprylate concentration in the fermentation broth (420 mM C; 7.57 g L^−1^). Given that methane production remained low at near-neutral and neutral pH, we theorized that the high concentration of undissociated *n*-caprylic acid (5.71 mM C) inhibited methanogenesis. We then demonstrated such an inhibitory effect at neutral pH in: **(1)** microcosm experiments; and **(2)** the continuous bioreactor by adding methanogenic sludge. Furthermore, 16S rRNA gene sequencing analysis revealed that near-neutral and neutral pH led to more diverse microbial communities than at mildly-acidic pH. For the first time, we report predominant *n*-caprylate production in a microbiome at near-neutral and neutral pH conditions where methanogenesis was controlled by the inhibitory effects of undissociated *n*-caprylic acid. At the same time, extraction of this species occurred even at near-neutral and neutral pH.

## 1. INTRODUCTION

Carbon-chain elongation *via* anaerobic fermentation has been referred to as a sustainable biotechnology production platform to convert carbon-rich organic waste streams into valuable industrial commodities and biofuels. Medium-chain carboxylates (MCCs) have chain lengths between 6 and 12 carbon atoms (C6-C12) and are industrial commodities. The relevance of MCCs lies in their various industrial applications, including their use as antimicrobials [1,2], additives for livestock feed [3], and precursors of liquid biofuel production [4–6]. The anaerobic production of MCCs from carbohydrate-rich substrates consists of two main metabolic modes. First, the substrate is converted *via* primary fermentation into short-chain carboxylates (SCCs, chain length: 2-5 carbons), following the classic anaerobic pathways Embden-Meyerhof-Parnas fermentation, lactic-acid fermentation, and ethanol fermentation [7]. Then, SCCs are further converted *via* secondary fermentation by reactor microbiomes into MCCs through the reverse β-oxidation and the fatty-acid biosynthesis pathways [8]. The microbiome uses some fraction of the substrate to produce energy and acetate for these processes.

In contrast, the rest of the substrate is channeled into acetyl-CoA or malonyl-ACP and sequentially elongates short-chain electron acceptors such as acetate. Chain-elongating microbiomes obtain additional energy by coupling the reverse β-oxidation with the formation of a cross-membrane proton gradient that ultimately translates into more ATP [9–11]. In the end, a mixture of MCCs, consisting mainly of *n*-caproate (C6), *n*-heptanoate (C7), and *n*-caprylate (C8), may be obtained when proper operating conditions and substrate quality is maintained. Industry prefers *n*-caprylate to the other MCCs due to its higher market value, reduced smell, and ease of separation [12].

The pH is critical among the multiple factors affecting microbial pathways, including reverse β-oxidation and fatty-acid biosynthesis. The pH controls the ratio of dissociated to undissociated carboxylates in a microbiome. Thus, as the pH becomes more acidic, the concentration of undissociated carboxylic acids increases. A mildly acidic pH (pH ∼ 5-6) is often preferred in chain-elongation microbiomes due to the desired inhibitory effect of a low pH on methanogenic activity [13–15], which is a pertinent concern because it can easily redirect the metabolic fluxes towards methane production. Acetoclastic methanogenesis leads to acetate consumption, reducing its availability for chain elongation. On the other hand, hydrogenotrophic methanogenesis could lower the partial pressure of H_2_ (below 10^−2^ atm), which leads to excessive ethanol oxidation, thereby depriving chain-elongation of its primary electron donor. Mildly acidic conditions also enhance the performance of MCC pertraction because the process is driven by the concentration gradient of undissociated carboxylic acid between the fermentation broth (pH: 5-6) and the extraction solution (pH: 9-11) [14,16–19].

Nevertheless, the transport rate of carboxylates across microbial membranes slows down at mildly acidic pH since it relies upon the concentration gradient of the undissociated forms [20–22]. In the cell, the accumulation of carboxylates leads to end-product feedback inhibition of reverse β-oxidation and disrupts protein synthesis [23–25]. Moreover, the increased concentration of undissociated carboxylic acids in the fermentation broth can also disrupt cell-membrane function by interfering with intracellular pH regulation [26,27].

Previously, we have operated a 5-L anaerobic sequencing batch reactor (ASBR) fed with corn beer from the corn-kernel-to-ethanol industry as the substrate for more than 1500 days at mesophilic conditions and a pH of 5.5. Volumetric production rates of *n*-caproate up to 180 mmol C L^−1^ d^−1^ (0.14 g L^−1^ h^−1^) were achieved [17,28]. Between days 552 and 1510, we operated the bioreactor in a low maintenance program; it turned out that the bioreactor had been unexpectedly operating at near-neutral pH (∼ 6.7) rather than 5.5 due to human error. Interestingly, at such conditions, high concentrations of *n*-caprylate were found both in the fermentation broth (>400 mmol C L^−1^) and the extraction solution (>50% of total MCCs) with nominal volumetric methane production rates. Our first theory was that the unmonitored pH drift to near-neutral promoted a shift in the chain-elongation metabolism toward the production of carboxylates of longer carbon chains (*i*.*e*., *n*-caprylate). This pH drift resulted in the maintenance of sufficiently low concentrations of undissociated carboxylic acids (mainly *n*-caprylic acid and *n*-caproic acid), ensuring its production while methanogens remained inhibited. A second theory was that the changes in MCC production resulted from the difference in the microbiome composition.

Therefore, the main objective of this study was to evaluate the effect of pH on chain-elongation performance and microbial community composition. Another objective was to investigate the mechanism of methanogen inhibition by undissociated *n*-caprylic acid at neutral pH (7) in an open culture system. To achieve these objectives, we continued operating the bioreactor that was fed corn beer and subjected to different pH values (from 5.5 to 7.0). The corn beer contained mainly ethanol, which served as the primary electron donor and carbon source. To investigate the mechanism of methanogen inhibition, we performed batch assays at different pH levels and undissociated *n*-caprylic acid concentrations.

## 2. MATERIALS AND METHODS

### 2.1 Inoculum and substrate

Initially, the >2100-day operating chain-elongating bioreactor was inoculated and started up with a natural microbiome from two sources: sheep rumen and a thermophilic anaerobic digester (Western Lake Superior Sanitary District, Duluth, MN) [14,17,28]. At the beginning of our present study, the bioreactor had been operating for 1510 days at different hydraulic retention times (HRT) in the range of 8-20 days and different loading rates in the range of 1.7-10.7 g COD L^−1^ d^−1^ [14,17,28]. Thus, the microbial community was well adapted to chain-elongation conditions. For the inhibition assays (batch experiments and Period V of the continuous bioreactor), we obtained the methanogenic inoculum from the mesophilic digester at a municipal wastewater treatment plant (Ithaca Wastewater Treatment Plant, Ithaca, NY), which treated a mixture of primary sludge and waste activated sludge. For all experiments, corn beer was used as substrate (Western New York Energy Co.), containing principally ethanol at a concentration of 122 g L^−1^.

### 2.2 Bioreactor setup and operation

We used an ASBR that we equipped with an in-line pertraction system, as described before (**Fig. S1-S2**) [17,28]. The experiments for the present study took place from Day 1510 onwards (from now on referred to as Day 0). The operating conditions for the ASBR consisted of 48-h cycles. We automatically controlled the temperature at 30 ± 1°C, while the pH was automatically controlled at values in the range of 5.5-7.0 by adding 5 M NaOH during mixing [14,17,28]. The bioreactor operating conditions consisted of five experimental periods during which we evaluated different pH levels, ranging from 5.5 to 7.0 (**Table 1**). During Period I (0-46 days), the bioreactor was operated at a pH of 6.75 (*i*.*e*., a similar pH that had been used for several years). We decreased the pH to 6.25 and 5.5 during Period II (48-182 days) and Period III (184-252 days), respectively. Afterward, the pH was increased from 5.5 to 6.0 during Period IV (254-316 days) and 7.0 during Period V (316-590 days) (**Table S1**). During Period V (526-590 days), we added methanogenic sludge to the bioreactor at a rate of 25 mL d^−1^. The bioreactor operated at an organic loading rate of 6.93 g COD L^−1^ d^−1^ and an HRT of 15 days for all experimental periods, including adding methanogenic sludge by displacing some corn beer.

**Table 1.** Bioreactor process performance during the five operating periods.

| Experimental phase | pH | Time (days) | Biomass sample (days) | OLR (g COD L<sup>-1</sup> d<sup>-1</sup>) | ELR (mmol C L<sup>-1</sup> d<sup>-1</sup>) | HRT (days) |
| --- | --- | --- | --- | --- | --- | --- |
| Period I | 6.75 | 0-46 | 28 | 6.93 | 79.66 | 15 |
| Period II | 6.25 | 48-182 | 112 | 6.93 | 79.66 | 15 |
| Period III | 5.50 | 184-252 | 212 | 6.93 | 79.66 | 15 |
| Period IV | 6.00 | 254-316 | 308 | 6.93 | 79.66 | 15 |
| Period V | 7.00 | 316-590 | 378 | 6.93 | 79.66 | 15 |

OLR: Organic Loading Rate; ELR: Ethanol Loading Rate.

### 2.3 Extraction system

As reported elsewhere, a pertraction system was used to avoid microbiome inhibition due to MCC accumulation [14,28]. For this purpose, two hollow-fiber membrane contactors *in series* (4 × 13 Extra-Flow, Liqui-Cel, USA) were connected to the bioreactor and the reservoir of extracted MCC solution (**Fig. S1**). Before entering the first contactor (*i*.*e*., forward contactor), the fermentation broth was drawn through a filter (FXWSC, GE Appliance, USA) to prevent membrane fouling. The filtered fermentation broth flowed through the forward contactor at a flow rate of 30 mL min^−1^. A hydrophobic solvent consisting of mineral oil (Sigma-Aldrich, USA) and 3% tri-*n*-octylphosphineoxide (Sigma-Aldrich, USA) was recirculated at a flow rate of 20 mL min^− 1^ between the forward and backward contactors. An extraction solution (pH 9.4 ± 0.2) continuously circulated through the back-membrane contactor with a flow rate of 25 mL min^−1^; this served as the final reservoir for produced MCCs. Each of the membrane contactors had a cross-sectional area of 8.10 m^2^.

### 2.4 Methanogenesis inhibition batch assays

We utilized 250-mL screw-cap bottles (working volume: 150 mL) for the methanogenesis inhibition batch assays. The active volume included: basal medium, N-N-bis(2-hydroxyethyl)-2-aminoethanesulfonic acid (BES), inoculum, and *n*-caprylate (**Table S4)** [29]. The headspace was flushed with pure N_2_ gas to displace oxygen and promote anaerobic conditions. We incubated the prepared bottles in triplicate at a temperature of 30°C.

### 2.5 Analytical methods and calculations

We sampled the fermentation broth and the extraction solution daily. The concentrations of ethanol, acetone, butanol, and carboxylic acids (C2 to C8) were analyzed by gas chromatography (GC) [13]. The soluble chemical oxygen demand (COD_s_), total chemical oxygen demand (COD_t_), ammonium, total solids (TS), and volatile solids (VS) concentrations were measured every week according to standard methods [30]. The biogas production was quantified using a liquid displacement device [17,28]. We determined the biogas composition by measuring hydrogen (H_2_), methane (CH_4_), nitrogen (N_2_), and carbon dioxide (CO_2_) through GC, as reported elsewhere [13]. The volumetric production rates of MCCs (mmol C L^−1^ day^−1^) were calculated by including both the carboxylic acids in the fermentation broth and the extraction solution.

### 2.6 Biomass sampling, sequencing, and microbiome analysis

We obtained five biomass samples from the bioreactor fermentation broth on Days 28, 112, 212, 308, and 378. The liquid samples were centrifuged in 2-mL Eppendorf tubes at 14,000 rpm for 5 min to separate biomass (the liquid supernatant was discarded). The biomass samples were stored at −80°C until further processing. Four biomass samples from a previous study with the same bioreactor were taken from the −80°C freezer (Days −1406; −991, −564, and −488 from Day zero). Genomic DNA was extracted from biomass samples using the DNeasy PowerSoil HTP 96 Kit (Qiagen, Germany), following the protocol detailed elsewhere [31]. The DNA samples were sequenced according to the protocol described by the Earth Microbiome Project [32]. In brief, the V4 regions of the 16S rRNA gene were amplified with the primers 515F (Parada) (5’-GTGYCAGCMGCCGCGGTAA) and 806R (Apprill) (5’-GGACTACNVGGGTWTCTAAT) that were fused with Illumina adapters and barcodes. The resulting amplicons were purified with the MoBio UltraClean PCR Clean-Up Kit (Qiagen, Germany). The amplicon library was sequenced using the Illumina MiSeq (1×150 bp) platform. The downstream sequence processing was performed using Quantitative Insights into Microbial Ecology (QIIME) software [33] following the pipeline described elsewhere [32,34]. The analysis included the quality filter, chimera check, and taxonomy assignment using the SILVA 16S rRNA gene database (132 release) as reference. The sequencing data were grouped into single variant reads (SVR) classified taxonomically to the genus level and expressed as relative abundance.

## 3. RESULTS AND DISCUSSION

### 3.1 A near-neutral pH level induced high *n*-caprylate production rates

We operated an ASBR (with continuous MCC extraction) at five different pH values (5.5-7.0) by adding different amounts of NaOH for a total of 590 days (**Fig. S3**). The maximum rate of volumetric *n*-caprylate production of 75.6 ± 0.6 mmol C L^−1^ d^−1^ (0.06 g L^−1^ h^−1^) and the highest concentration of *n*-caprylate in the fermentation broth of 420 mM C (7.57 g L^−1^) were observed during Period I (pH: 6.75) (**Table 2, Fig. 1A-B**). Distinctly, the pH affected the ratio of *n*-caprylate to *n*-caproate (C8 to C6) in the extraction solution (**Fig. 2A**). The near-neutral pH resulted in a lower *n*-caprylate extraction efficiency of 63% compared to 96% at a pH of 5.5 (**Table 2, Fig. 2B**), resulting in a relatively high *n*-caprylate concentration in the fermentation broth.

**Table 2.**
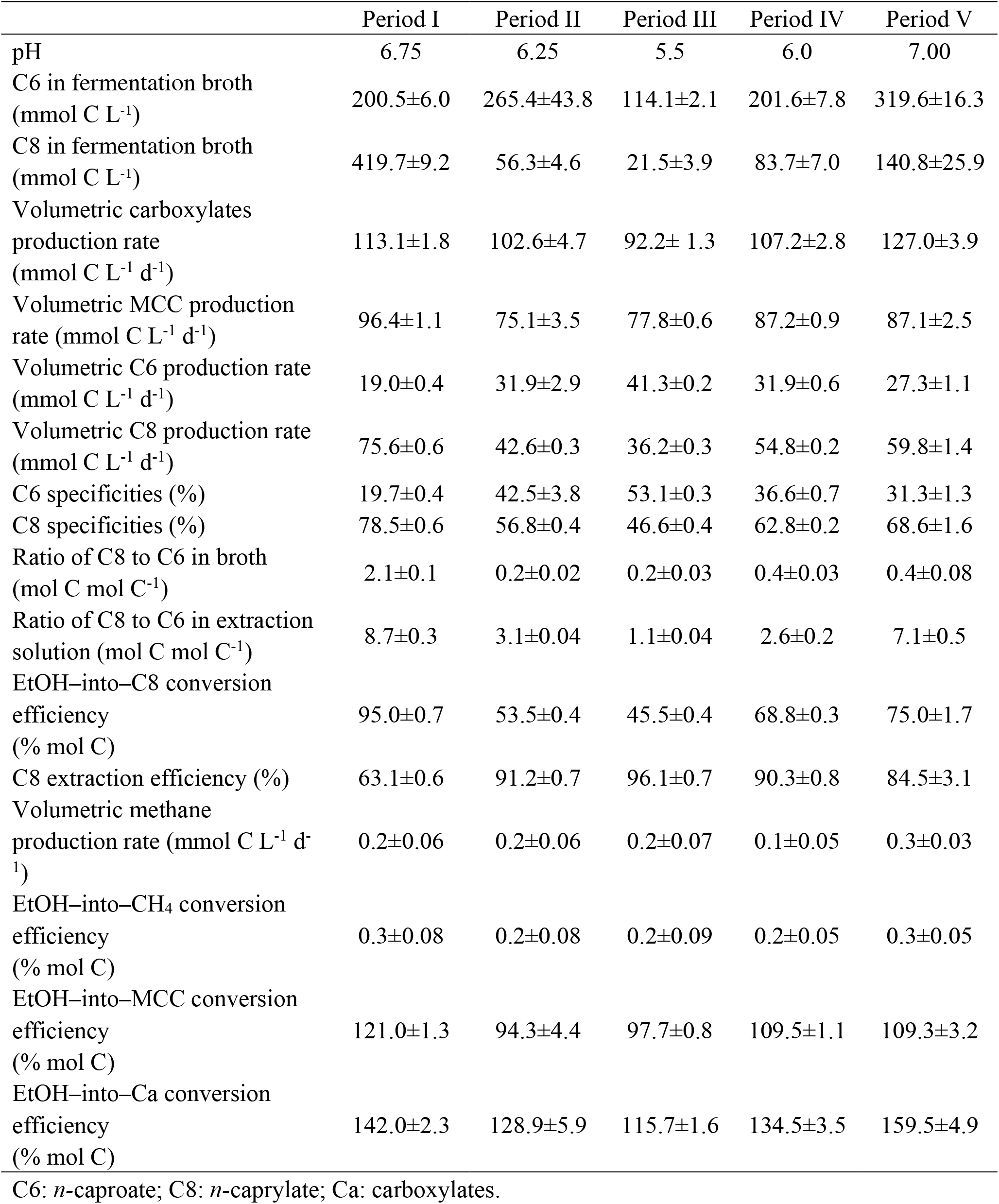
Summary of results data for the five operating periods: volumetric production rate, C8–to–C6 ratio in the extraction solution, conversion efficiency, concentration, and extraction efficiency.

|  | Period I | Period II | Period III | Period IV | Period V |
| --- | --- | --- | --- | --- | --- |
| pH | 6.75 | 6.25 | 5.5 | 6.0 | 7.00 |
| C6 in fermentation broth<br>(mmol C L<sup>-1</sup>) | 200.5±6.0 | 265.4±43.8 | 114.1±2.1 | 201.6±7.8 | 319.6±16.3 |
| C8 in fermentation broth<br>(mmol C L<sup>-1</sup>) | 419.7±9.2 | 56.3±4.6 | 21.5±3.9 | 83.7±7.0 | 140.8±25.9 |
| Volumetric carboxylates<br>production rate<br>(mmol C L<sup>-1</sup> d<sup>-1</sup>) | 113.1±1.8 | 102.6±4.7 | 92.2± 1.3 | 107.2±2.8 | 127.0±3.9 |
| Volumetric MCC production<br>rate (mmol C L<sup>-1</sup> d<sup>-1</sup>) | 96.4±1.1 | 75.1±3.5 | 77.8±0.6 | 87.2±0.9 | 87.1±2.5 |
| Volumetric C6 production rate<br>(mmol C L<sup>-1</sup> d<sup>-1</sup>) | 19.0±0.4 | 31.9±2.9 | 41.3±0.2 | 31.9±0.6 | 27.3±1.1 |
| Volumetric C8 production rate<br>(mmol C L<sup>-1</sup> d<sup>-1</sup>) | 75.6±0.6 | 42.6±0.3 | 36.2±0.3 | 54.8±0.2 | 59.8±1.4 |
| C6 specificities (%) | 19.7±0.4 | 42.5±3.8 | 53.1±0.3 | 36.6±0.7 | 31.3±1.3 |
| C8 specificities (%) | 78.5±0.6 | 56.8±0.4 | 46.6±0.4 | 62.8±0.2 | 68.6±1.6 |
| Ratio of C8 to C6 in broth<br>(mol C mol C<sup>-1</sup>) | 2.1±0.1 | 0.2±0.02 | 0.2±0.03 | 0.4±0.03 | 0.4±0.08 |
| Ratio of C8 to C6 in extraction<br>solution (mol C mol C<sup>-1</sup>) | 8.7±0.3 | 3.1±0.04 | 1.1±0.04 | 2.6±0.2 | 7.1±0.5 |
| EtOH-into-C8 conversion<br>efficiency<br>(% mol C) | 95.0±0.7 | 53.5±0.4 | 45.5±0.4 | 68.8±0.3 | 75.0±1.7 |
| C8 extraction efficiency (%) | 63.1±0.6 | 91.2±0.7 | 96.1±0.7 | 90.3±0.8 | 84.5±3.1 |
| Volumetric methane<br>production rate (mmol C L<sup>-1</sup> d<sup>-1</sup>) | 0.2±0.06 | 0.2±0.06 | 0.2±0.07 | 0.1±0.05 | 0.3±0.03 |
| EtOH-into-CH<sub>4</sub> conversion<br>efficiency<br>(% mol C) | 0.3±0.08 | 0.2±0.08 | 0.2±0.09 | 0.2±0.05 | 0.3±0.05 |
| EtOH-into-MCC conversion<br>efficiency<br>(% mol C) | 121.0±1.3 | 94.3±4.4 | 97.7±0.8 | 109.5±1.1 | 109.3±3.2 |
| EtOH-into-Ca conversion<br>efficiency<br>(% mol C) | 142.0±2.3 | 128.9±5.9 | 115.7±1.6 | 134.5±3.5 | 159.5±4.9 |

C6: *n*-caproate; C8: *n*-caprylate; Ca: carboxylates.

**Fig. 1.**
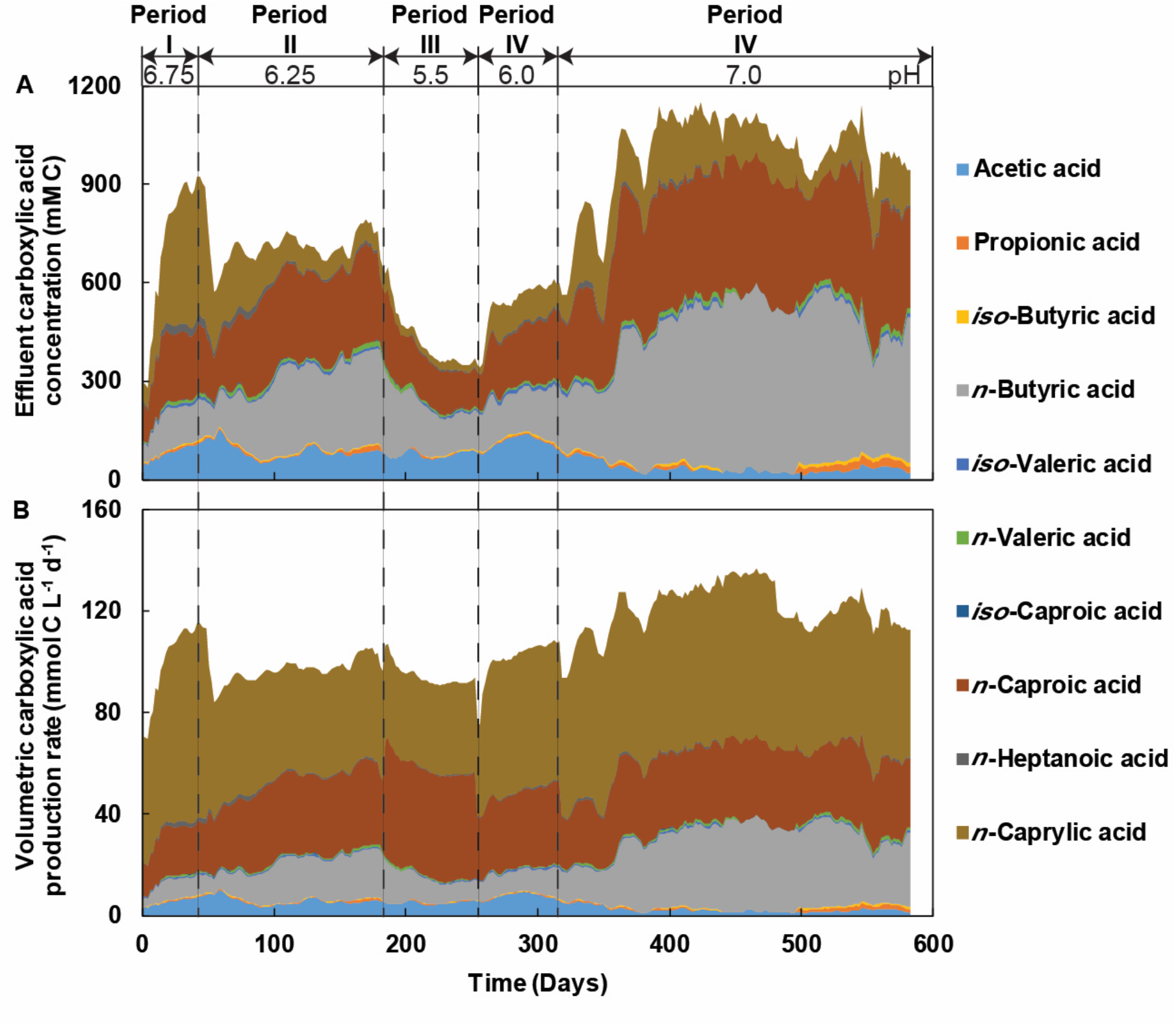
Performance, fermentation broth concentration, and volumetric production rate during Periods I to V. The fermentation broth concentration of carboxylic acids (A) and volumetric production rates of carboxylic acids (B) for all periods.

**Fig. 2.**
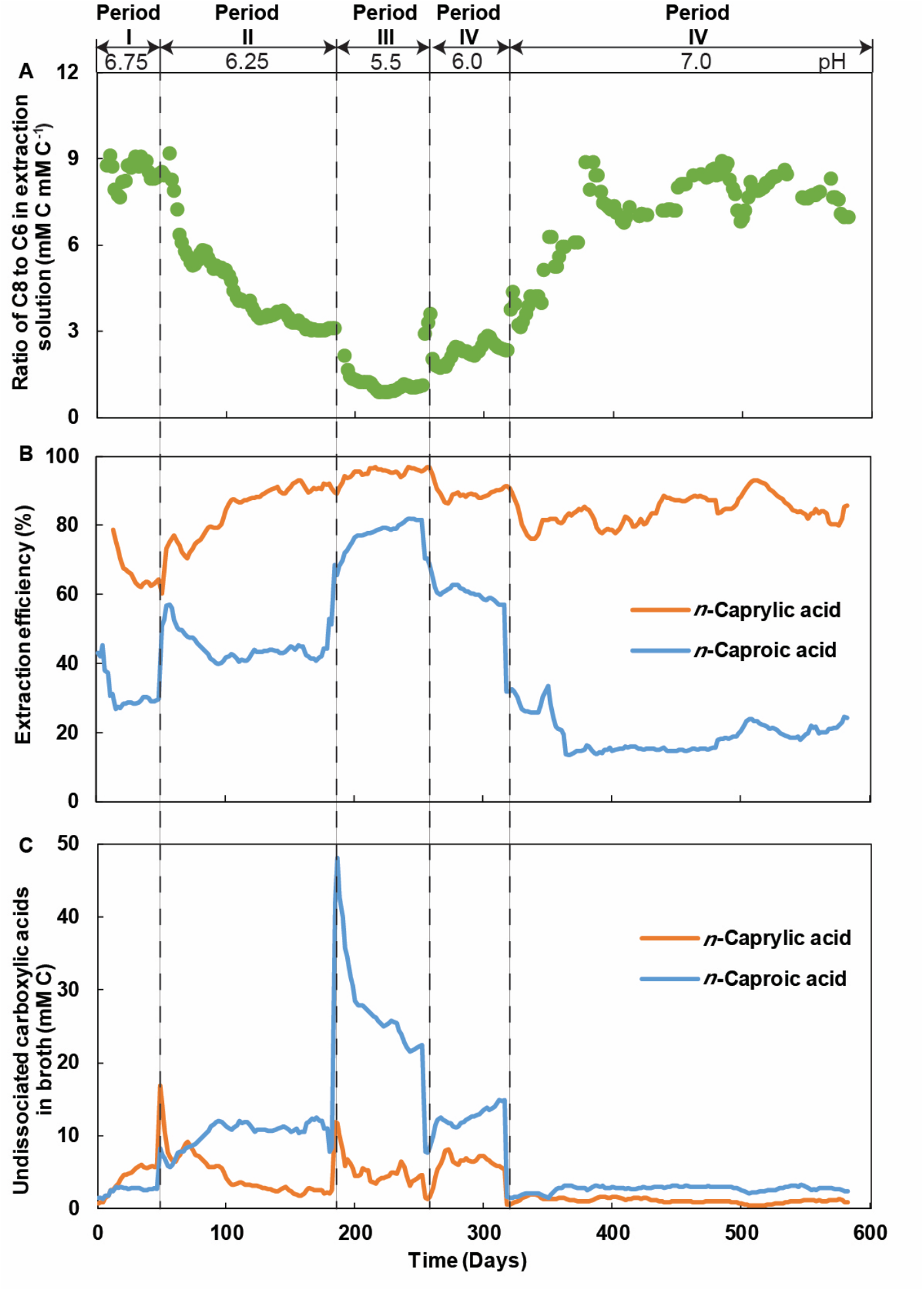
The ratio of *n*-caprylate to *n*-caproate (C8 to C6) in the extraction solution (A), extraction efficiency (B), and undissociated carboxylic acids in the fermentation broth based on the global pKa of ∼4.9 (C) during the operating periods.

Therefore, the concentration of *n*-caprylate in the fermentation broth at a pH of 6.75 was approximately eightfold higher than the maximum *n*-caprylate concentration of 50 mM C (0.9 g L^−1^) [35] that was reported thus far (**Fig. S4A**). Moreover, at such conditions, we estimated, with the global pKa of 4.9 (**Fig. S5**), a maximum concentration of undissociated *n*-caprylic acid of 5.71 mM C (0.10 g L^−1^), which was considerably higher than those reported previously as well. Our study also reached the highest ethanol-to-*n*-caprylate conversion efficiency thus far (**Fig. S4B**), indicating the excellent removal of ethanol (**Fig. S6)**. The high concentration of undissociated *n*-caprylic acid found in our system also suggested that the carboxylate feedback inhibition, which often completely inhibits the chain-elongation processes, was not a relevant problem in our system. Instead, our results indicate a selection of chain-elongating microbes that could survive under high concentrations of undissociated *n*-caprylic acid [36] (further discussion in Sections 3.3 and 3.4).

During Period I (pH of 6.75), we also observed the maximum rate of the volumetric total MCC production (96.4 ± 1.1 mmol C L^−1^ d^−1^), *n*-caprylate specificity (78.50%), and *n*-caprylate–to–*n*-caproate (C8–to–C6) ratio (3.99 mol C mol C^−1^) (**Table 2**). During Period II, the pH was decreased from 6.75 to 6.25, which resulted in a slower rate of volumetric *n*-caprylate production (42.6 mmol C L^−1^ d^−1^) and reduced *n*-caprylate specificity (46.6%) (**Table 2**). A further decrease in the pH to 5.50 (Period III) led to the minimum rate of volumetric *n*-caprylate production (36.2 mmol C L^−1^ d^−1^). In contrast, the average volumetric *n*-caproate production rate increased from 18.95 to 31.90 and then to 41.31 mmol C L^−1^ d^−1^ when the pH was decreased from 6.75 (Period I) to 6.25 (Period II) and then to 5.5 (Period III), respectively. Such a trend of volumetric *n*-caproate production rate was accompanied by an increase in *n*-caproate specificity from 19.7% during Period I to 53.1% during Period III. The overall volumetric MCC production rates, which included both *n*-caproate and *n*-caprylate, had changed only somewhat between the periods, with a maximum rate of 96 mmol C L^−1^ d^−1^ at a pH of 6.75, and a minimum rate of 75 mmol C L^−1^ d^−1^ at a pH of 6.25 (**Table 2**).

These results suggest that the decrease of pH minimized the *n*-caprylate conversion rate due to an intracellular *n*-caprylate accumulation, which hindered the conversion of *n*-caproate to *n*-caprylate (end-product feedback inhibition). In turn, the intracellular accumulation of *n*-caprylate likely resulted from a high share of extracellular undissociated *n*-caprylic acid, which interfered with free carboxylic acid diffusion from the cytoplasm (generally at a neutral pH) to the fermentation broth [25]. On the contrary, under the decreased pH conditions, the production of *n*-caproate was stimulated by the lower toxicity of *n*-caproate compared to *n*-caprylate. In addition, we observed the most promising hydrolysis efficiencies at the pH values of 5.5 and 6.0 (Period II and IV) with the lowest effluent COD concentrations (**Fig. S7A**), the highest effluent ammonium concentrations (**Fig. S7B**), and the lowest effluent solids concentrations (**Fig. S7C**) compared to the other periods.

To confirm that *n*-caprylate production was favored at pH > 5.5, we increased the pH from 5.5 to 7.0 in two steps during Periods IV and V. During Period IV, the pH was set to 6.0, resulting in an increase of the volumetric *n*-caprylate production rate to 54.8 mmol C L^−1^ d^−1^, while the *n*-caprylate concentration increased to 83.7 mM C. Similarly, the *n*-caprylate specificity changed from 46.6% to 62.8%. The positive effect of increasing the pH on *n*-caprylate production was confirmed when further increasing the pH to 7.0 during Period V. At such conditions, the *n*-caprylate production rate and concentration reached 59.8 mmol C L^−1^ d^−1^ and 140.8 mM C, respectively. The considerably higher concentration of *n*-caprylate at a pH of 7.0 than 5.5 also showed that the extraction efficiency of *n*-caprylate was reduced at the higher pH (from 96% to 85% in **Table 2, Fig. 2B**). Thus, the *n*-caprylate concentration increased due to a higher production rate and lower extraction efficiencies at the neutral pH.

The change in pH from 5.5 to 6.0 and then to 7.0 led to an increase of *n*-caprylate specificity from 46.6 to 62.8 and then to 68.6%, respectively. Even though the neutral conditions (pH 7.0) drove the system back to relatively high *n*-caprylate production rates and specificities, the performance was slightly lower than that observed at a pH of 6.75. Such a difference could be explained by possible changes in the microbial communities (see Section 3.4). The relatively inferior performance at a pH of 7.0 than 6.75 was accompanied by a considerably higher concentration of *n*-butyrate (Period I: 117.5 mM C; Period V: 521.4 mM C in **Fig. 1A**), which suggested that chain elongation was limited somewhat at the neutral pH level.

Overall, we found that the pH of 6.75 (Period I) was optimal for *n*-caprylate production in a performance situation that *n*-caprylate was produced sufficiently to inhibit methanogens. Although low pH has been referred to as a favorable factor for chain elongation [37,38], our results showed that near-neutral pH supported the production of *n*-caprylate. This result was unexpected given the conventional assumption that chain elongation performance should drop at near-neutral pH due to the lower extraction efficiencies and the appearance of competing metabolisms such as methanogenesis. We hypothesize that the relatively high pH promoted the process of chain elongation by decreasing the concentration of undissociated carboxylic acids and avoiding intracellular product accumulation. At such conditions, the *n*-caprylate accumulated in the fermentation broth at concentrations that probably avoided the rise of methanogenic archaea and other competing microbes.

An explanation for this unexpected result may be that the concentrations and ratios of dissociated and undissociated *n*-caprylic acid can be affected by the formation of micelles, which are favored at high pH, as discussed by Urban and Harnisch [39]. On the one hand, the appearance of micelles means establishing a biphasic system, which decreases the concentration of undissociated *n*-caprylic acid in the fermentation broth due to partitioning between the aqueous and organic phases. The decrease in undissociated *n*-caprylic acid concentration in the aqueous phase should effectively lower its toxicity. On the other hand, the formation of micelles could cause a localized increase in pKa at the micelle surface due to the coincident arrangement of polar carboxyl groups from the carboxylic acid molecules. Ultimately, a locally higher pKa than the global pKa (**Fig. S5**) translates into higher concentrations of undissociated *n*-caprylate than would be expected at any given pH. For mildly acidic pH values, this would explain the lower performance observed in our bioreactor at the pH of 5.5 due to enhanced toxicity. For near-neutral or neutral pH values, on the other hand, this would explain why pertraction still worked for undissociated *n*-caprylic acid. The superior extraction efficiency for *n*-caprylate *vs. n*-caproate at neutral pH values proves that micelle formation plays a pertinent role for pertraction; without it, the pertaction efficiency for *n*-caprylate should have been as similarly low as for *n*-caproate (>80% *vs*. <20% at pH of 7.0 in **Fig. 2B**).

### 3.2 A pH-dependent pertraction system allowed high-rate *n*-caprylate production

We operated a continuous product extraction system to overcome product inhibition and increase MCC production. This pertraction system selectively extracted MCCs from the fermentation broth through the forward membrane contactor into a hydrophobic solvent and then through the backward membrane contactor into the extraction solution [13,15,17,28]. In such a system, the hydrophobic solvent selects longer undissociated MCCs due to their more hydrophobic nature. With micelle formation for only undissociated *n*-caprylic acid at near-neutral and neutral pH, this selection effect of longer undissociated MCCs was even more pronounced because of the superior extraction of C8 *vs*. C6 at near-neutral or neutral pH (**Fig. 2B**). For undissociated *n*-caproic acid, the concentration at near-neutral and neutral pH conditions without micelle formation is so low (*e*.*g*., 0.77% of the total *n*-caproate in **Fig. S5**) that the extraction rate for *n*-caproate would be relatively slow, selecting for *n*-caprylate extraction and production.

Indeed, we observed the highest C8–to–C6 ratio of 2.1 (mM C mM C^−1^ in **Table 2**) in the fermentation broth during Period I with a pH of 6.75, leading to the highest C8–to–C6 ratio of 8.66 (mM C mM C^−1^ in **Table 2, Fig. 2A**) in the extraction solution. A decrease of the pH from 6.75 to 6.25 (Periods I to II) resulted in a considerable reduction in C8– to–C6 ratios of 0.21 and 3.09 (mol C mol C^−1^) in the fermentation broth and extraction solutions, respectively (**Table 2, Fig. 2A**). A further decrease of pH from 6.25 to 5.5 (Periods II to III) led to an additional reduction of the C8–to–C6 ratios in the fermentation broth and extraction solution of 0.19 and 1.12, respectively (**Table 2, Fig. 2A**). We observed consistent increases in the C8–to–C6 ratios when we increased the pH. From Period IV to VI, the pH increased from 5.5 to 6.0 and 7.0. As anticipated from the previous periods, the increases of pH led to increases in the C8–to–C6 ratio in both the fermentation broth and extraction solution (**Fig. 2A, Table 2**).

### 3.3 High concentrations of MCCs inhibited the methanogenic activity

Chain elongation is an anaerobic technology for organic waste treatment that is feasible in open-culture conditions [40]. This feature confers a lower cost compared to pure-culture systems. However, in open cultures with long retention times, organic substrates are readily converted into methane [11]. Therefore, targeted strategies to circumvent methane formation are needed to ensure an efficient MCC production process. Previous approaches in this regard have included: **(1)** operating at neutral pH with methanogen inhibitors (*e*.*g*., 2-bromoethanosulfonic acid (BrES) [40]); **(2)** operating at neutral pH (6.5-7.0) and a relatively short HRT (4-17 h) [41,42], and **(3)** operating at mildly acidic pH levels (*i*.*e*., 5.0-5.5) [12,13,43]. Here, we observed low volumetric methane production rates of 0.1-0.3 mmol C L^−1^ d^−1^ throughout the range of pH conditions (from 5.5 to 7.0), which equate to losses of 0.2-0.3% relative to total ethanol inputs (**Table 2, Fig. 3A**). These results indicate that the methanogenic activity was inhibited at all pH conditions tested (including near-neutral and neutral pH) at a relatively long HRT (15 days) without implementing any additional strategy.

**Fig. 3.**
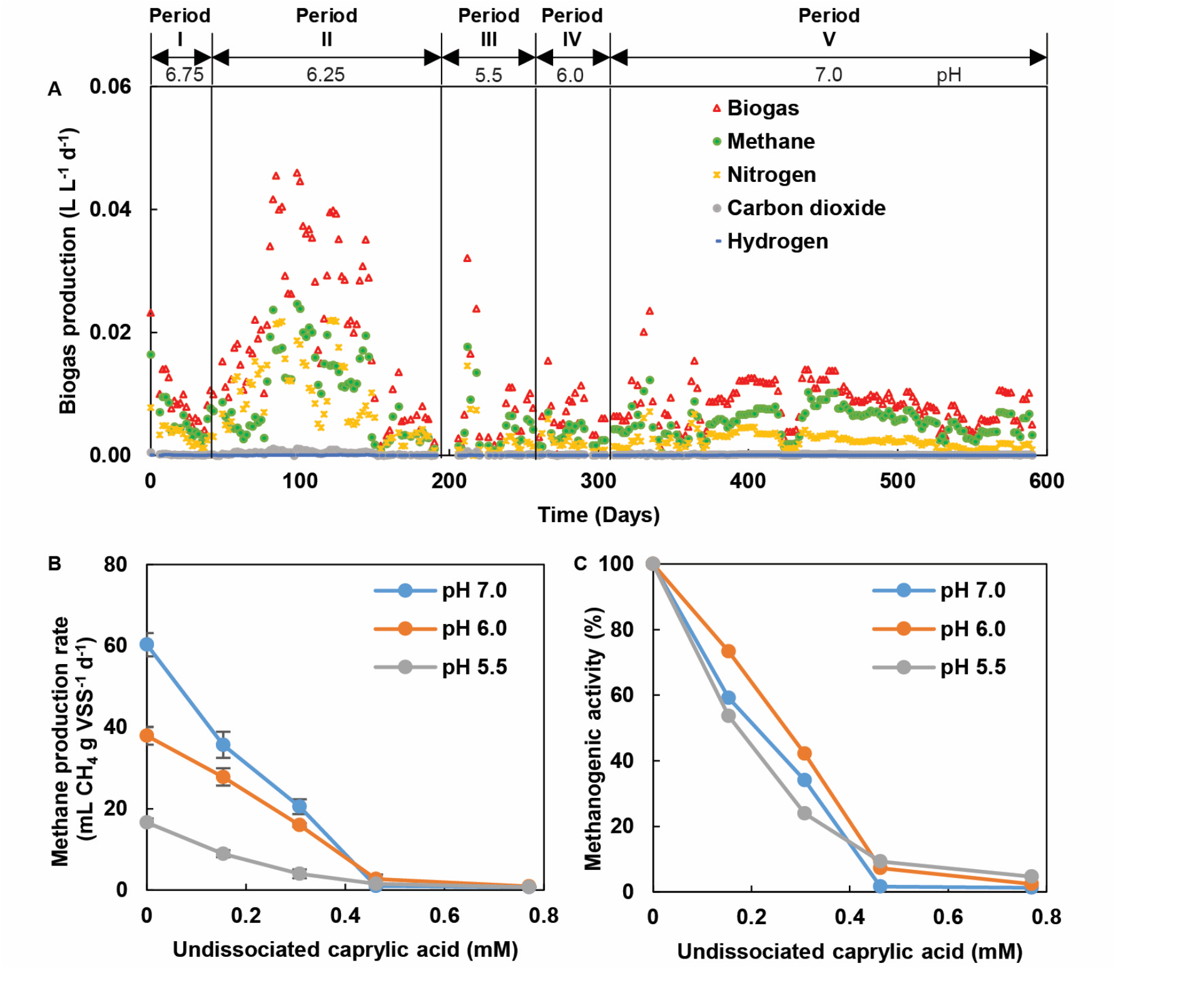
Biogas production during the operating periods and batch methanogen inhibition assays. (A) Biogas production data including methane, nitrogen, carbon dioxide, and hydrogen during all experimental periods. (B) Methane production with different pH and concentrations of undissociated *n*-caprylic acid under batch methanogen inhibition assays. (C) Methane activity with different pH and concentrations of undissociated *n*-caprylic acid under batch methanogen inhibition assays.

We theorized that the suppression of methanogenesis resulted from the high concentrations of MCCs, which seemed to inhibit methanogenic activity. Carboxylic acids have been widely used as food preservatives for decades due to their antimicrobial properties [44]. Different authors report that SCCs and MCCs can inhibit the growth of *Escherichia coli* [45–47], *Salmonella* spp. [47–49], *Saccharomyces cerevisiae* [50], *Shigella dysenteriae* [48], *Vibrio cholerae* [48], *Clostridium perfringens* [47], and eggs of the roundworm *Ascaris suum* [1,51]. Many studies have also shown that the toxicity of carboxylates increases with increasing carbon-chain length (up to 12 C) [44–46,51,52]. Koster et al. [53] already showed that *n*-caprylate, *n*-decanoate, and other long-chain carboxylic acids could inhibit acetoclastic methanogenic activity at near-neutral pH.

To confirm that relatively high concentrations of *n*-caprylate inhibit methanogenesis, we investigated the methane production rate and methanogenic activity at different pH levels and concentrations of undissociated *n*-caprylic acid (**Fig. 3A-B**). Results showed that undissociated *n*-caprylic acid at 0.46 mM C caused more than 90% inhibition of methane production rate and methanogenic activity at all pH levels (5.5, 6.0, and 7.0) (**Fig. 3B-C**). The equivalent total *n*-caprylate (undissociated and dissociated) concentrations were 2.34 mM C, 6.41 mM C, and 59.9 mM C at the pH of 5.5, 6.0, and 7.0, respectively. Considering that the concentration of undissociated *n*-caprylic acid was well above 0.46 mM C at all conditions tested, it played a role in inhibiting methanogenesis.

We also further investigated the effectiveness of methanogenesis inhibition in the continuous bioreactor. For this purpose, from day 526 to day 590, we fed active methanogenic sludge at a rate of 25 mL d^−1^ at a neutral pH of 7.0. In agreement with the results of the batch experiments, methane production in the continuous bioreactor did not increase, remaining at 0.3 mmol C L^−1^ d^−1^ (**Fig. 3A**) with an *n*-caprylate concentration of 142 mM, which is well above concentrations considered inhibitory to methanogenesis. Methanogenesis was actively limited by *n*-caprylate, preventing the conversion of the chain-elongating system into a methanogenic anaerobic digestion system (**Fig. 3**).

### 3.4 Greater microbial diversity at high pH drove *n*-caprylate production

The 16S rRNA gene sequencing analysis showed that the pH of operation substantially affected the composition and diversity of microbial communities (**Fig. 4**). In particular, the bioreactor was dominated by microbes belonging to the genera *Caproiciproducens, Ruminococcus, Oscilibacter*, and *Methanobacterium*. Other sub-dominant bacteria belonging to the genera *Erysipelatoclostridium, Anaerosporobacter*, and *Actinomyces* were also present, especially during the periods of high volumetric *n*-caprylate production (**Fig. 4**). We also identified these microbes in samples taken from previous periods of operation (*i*.*e*., Day −1406 [pH 5.5], Day −991 [pH 5.5], Day −564 [pH 5.81], and Day −488 [pH 5.91] in **Fig. 4**).

**Fig. 4.**
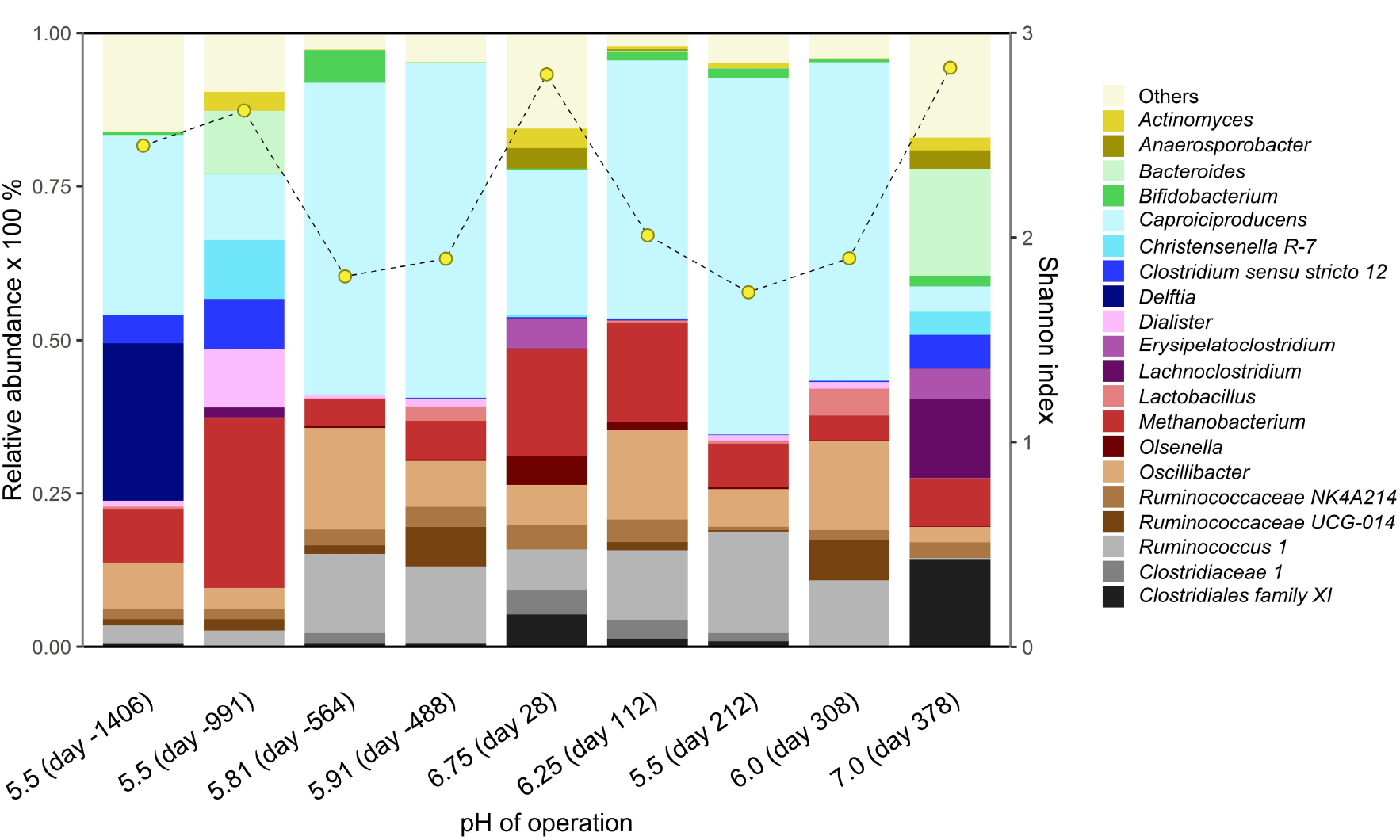
Composition and diversity of microbial communities involved in medium chain carboxylic acids production at different pH levels. The taxonomic level was cut off at the genus rank. The group denoted as Others, includes the unique variant reads with a cumulative abundance across samples lower than 5%. The dashed line corresponds to the Shannon diversity index.

For the dominant microbes that were present, we found that the relative abundance of *Caproiciproducens* spp. was strongly related to *n*-caproate production, which occurred at higher rates under mildly acidic conditions (pH: 5.5). Several species of this genus have been identified in *n*-caproate-producing microbiomes utilizing sugars as the electron donor at mildly-acidic conditions [54–56]. Similarly, we found that the presence of *Ruminococcaceae* genera (*Ruminococcus* 1, *Ruminococcaceae* N4A214, *Ruminococcaceae* UCG-014) was also related to the production of *n*-caproate at mildly-acidic conditions. This family of bacteria has also been reported in multiple chain-elongating bioreactors [56]. Both the *Caproiciproducens* and *Ruminococcaceae* genera were favored at mildly-acidic pH (54.9% and 15.6%, respectively), while their presence at near-neutral and neutral pH (≥ 6.75) was less dominant (<20.9 and < 5.9%, respectively).

Because subdominant species can also be key players in shaping metabolic pathways, the microbes (*Erysipelatoclostridium, Anaerosporobacter*, and *Actinomyces*) are worth considering. *Erysipelatoclostridium* belongs to the family of Erysipelatoclostridiaceae, which are gram-positive anaerobic bacteria that share phenotypic features with members of the Firmicutes phylum. Among other characteristics, this means that *Erysipelatoclostrium* (as well as other members of the Erysipelatoclostridiaceae) is an anaerobic fermentative bacterium that can ferment multiple substrates into lactate and other short-chain carboxylates [57]. Given that the relative abundance of *Erysipelatoclostridium* increased during periods characterized by higher rates of volumetric *n*-caprylate production (pH 6.75 and 7.0), it seems clear that *Erysipelatoclostridium* participated in either lactate-based chain elongation or the acidogenic phase of primary fermentation.

The genus *Actinomyces* coincided with high concentrations of *n*-caprylate. *Actinomyces* are lactate-producing facultative anaerobes commonly observed in chain-elongation systems [58–60]. Their role in the microbial community seems to be associated with lactate production; however, due to their facultative nature, they could also contribute by ensuring anaerobic conditions or through propionate production. Microbes belonging to the *Anaerosporobacter* genus were also found during periods of high *n*-caprylate production rates. *Anaerosporobacter* is anaerobic fermentative microbes whose main products are formate, acetate, and H_2_. Their role in the microbial community is not well defined. Our 16S rRNA gene sequencing analysis confirmed the presence of methanogens throughout the operating period. Still, as described previously, volumetric methane production rates remained below 0.4 mmol C L^−1^ d^−1^ (**Table 2**). Despite their low activity, we do not understand how methanogens persisted in the microbiome during this long period.

In general, the microbial community analysis showed that the pH affected the composition of the microbial community and its diversity. During the periods of high *n*-caprylate production, the microbial diversity as measured by the Shannon index, which quantifies both the richness and evenness of the community, reached maximum values of 2.79 and 2.83 in samples at pH of 7.0 and 6.75, respectively. On the contrary, the microbial community became less diverse as the pH decreased (Shannon index: 1.73 at pH 5.5). In summary, it seems that less diverse communities were involved in the pathways leading to *n*-caproate production, which we assume to be less complex compared to the pathways that are required for *n*-caprylate synthesis.

The literature has deliberated on the relationship between microbial diversity and process performance. However, few scientists have addressed this topic in their studies around chain elongation. Most microbial ecologists argue that microbial diversity is associated with more stable performance^17^. We did not observe problems due to instability; still, there was a clear association between microbial diversity and the microbiome’s function. A higher diversity correlated with the production of more complex compounds. The formation of these complex compounds probably requires the participation of a greater number of microbial species compared to the number needed to form simpler compounds. Nevertheless, many questions remain for future studies regarding the function of specific species within the chain elongation microbiome.

## 4. CONCLUSIONS

We investigated the effects of pH on the production and selectivity of *n*-caprylate in an ASBR microbiome with continuous pertraction. We found that mildly acidic pH (5-6) favored the *n*-caproate (C6) production, while near-neutral and neutral pH (6.75-7.0) steered the fermentation towards the production of *n*-caprylate. In particular, the pH of 6.75 led to a maximum volumetric *n*-caprylate production rate of 75.6 ± 0.6 mmol C L-1 d-1 (0.06 g L^−1^ h^−1^) with an *n*-caprylate concentration of 420 mM C (7.57 g L^−1^) in the fermentation broth. At such conditions, we estimated a concentration of undissociated *n*-caprylic acid of 5.71 mM C (0.10 g L^−1^), which is the highest reported up to date. We theorize that the high concentration of undissociated *n*-caprylic acid has an inhibitory effect on methanogenesis, which remained low at near-neutral and neutral pH. Moreover, the 16S rRNA gene sequencing analysis revealed that the microbial diversity was higher at near-neutral and neutral pH (6.75-7.0) than mildly acidic pH (5-6), suggesting that *n*-caprylate production requires the participation of a greater number of microbial species compared to the number needed to form simpler compounds. Overall, we report the predominant *n*-caprylate production in a microbiome at near-neutral and neutral pH conditions where methanogenesis is controlled by the inhibitory effects of undissociated n-caprylic acid. At the same time, extraction of this species occurs even at near-neutral and neutral pH, albeit at lower efficiencies than at mildly acidic pH.

## ACKNOWLEDGEMENTS

RPB acknowledges the support of the Alexander von Humboldt Foundation for the Georg Forster Research Fellowship. LA acknowledges support from the Alexander von Humboldt Foundation in the Alexander von Humboldt Professorship framework endowed by the Federal Ministry of Education and Research in Germany. The Earth Microbiome Project performed sample processing, sequencing, and core amplicon data analysis (www.earthmicrobiome.org), and all amplicon sequence data and metadata have been made public through the EMP data portal (qiita.microbio.me/emp). This work was supported by the U.S. Army Research Laboratory and the U.S. Army Research Office [grant number W911NF-12-1-0555]. We acknowledge the owners and operators of Western New York Energy for providing yeast-fermentation beer.

