## Supporting Information for "Near-neutral pH increased *n*-caprylate production in a microbiome with product inhibition of methanogenesis"

### Equations

#### Volumetric ethanol loading rate (mmol C L$^{-1}$ d$^{-1}$):

$$\frac{C_{EtOH} \times 2 \times f}{V} \qquad \text{(Eq. S1)}$$

Where:

$C_{EtOH}$ = concentration of ethanol in the fermentation broth, mM

f = effluent flow rate, L d$^{-1}$

V = the volume of reactor, L

#### Volumetric SCOD loading rate (g COD L$^{-1}$ d$^{-1}$):

$$\frac{SCOD \times f}{V} \qquad \text{(Eq. S2)}$$

Where:

SCOD = the concentration of soluble chemical oxygen demand in the influent, g COD L$^{-1}$

f = effluent flow rate, L d$^{-1}$

V = reactor volume, L

#### Specificities (%, mol C L$^{-1}$ d$^{-1}$):

$$\frac{\gamma_s}{\sum_{i=1}^{n} \gamma_i} \times 100 \qquad \text{(Eq. S3)}$$

Where:

$\gamma_s$ = production rate of a specific product, mmol C L$^{-1}$ d$^{-1}$

$\gamma_i$= production rate of MCCAs, mmol C L$^{-1}$ d$^{-1}$

#### Volumetric production rate (mmol C L$^{-1}$ d$^{-1}$):

$$\left[\frac{C_{e,n}V}{HRT} + \frac{(C_{b,n} - C_{b,n-1})V_b}{T_n - T_{n-1}}\right] \times \frac{1}{V} \qquad \text{(Eq. S4)}$$

Where:

$C_{e,n}$= concentration of the carboxylate in the fermentation broth on the day n, mM C

V = reactor volume, L

$HRT$ = hydraulic retention time on the day n, d

$C_{b,n}$, $C_{b,n-1}$ = concentrations of carboxylate in the stripping solution on the day n and n-1, mM C

$V_b$ = volume of the stripping solution on the day n, L

$T_n$ = the day n, d

#### Product-to-carboxylates production ratio (%, mol C L$^{-1}$ d$^{-1}$):

$$\frac{\gamma_s}{\sum_{i=1}^{n} \gamma_i} \times 100 \qquad \text{(Eq. S5)}$$

Where:

$\gamma_s$ = production rate of a specific product, mmol C $L^{-1}$ $d^{-1}$

$\gamma_i$= production rate of all carboxylates, mmol C $L^{-1}$ $d^{-1}$

#### Substrate-into-product conversion efficiency (%, mol C $L^{-1}$ $d^{-1}$):

$$\frac{\gamma_s}{\sum_{i=1}^{n} C_i \times f \times N_i / V} \quad \text{(Eq. S6)}$$

Where:

$\gamma_s$ = production rate of specific product, mmol C $L^{-1}$ $d^{-1}$

$C_i$ = concentration of specific substrate, mM C

f = effluent flow rate, L $d^{-1}$

$N_i$ = the number of carbons in a specific substrate

V = reactor volume, L

#### SCOD conversion efficiency (%, g COD)

$$\frac{\gamma_s}{SCOD_{LR}} \quad \text{(Eq. S7)}$$

Where:

$\gamma_s$ = production rate of a specific product, g COD $L^{-1}$ $d^{-1}$

$SCOD_{LR}$ = volumetric SCOD loading rate, g COD $L^{-1}$ $d^{-1}$

#### Molar percentage of MCCAs (%, mM):

$$\frac{C_i}{\sum_{i=1}^{n} C_i} \times 100 \quad \text{(Eq. S9)}$$

Where:

$C_i$ = concentration of carboxylate i, mM

### Figures

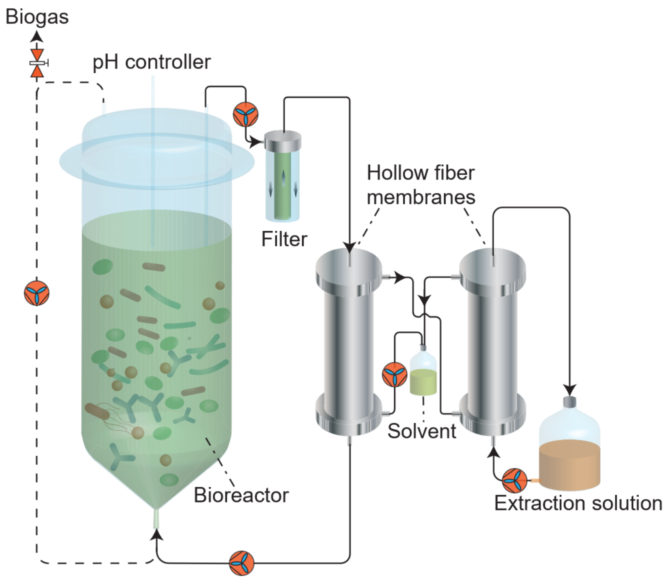

**Figure S1**. Schematic of the bioreactor and membrane-based liquid-liquid extraction system. Solid lines represent liquid streams; dashed lines represent gas streams. Feed and effluent were removed every other day and lines are not shown here (semi-batch operating conditions).

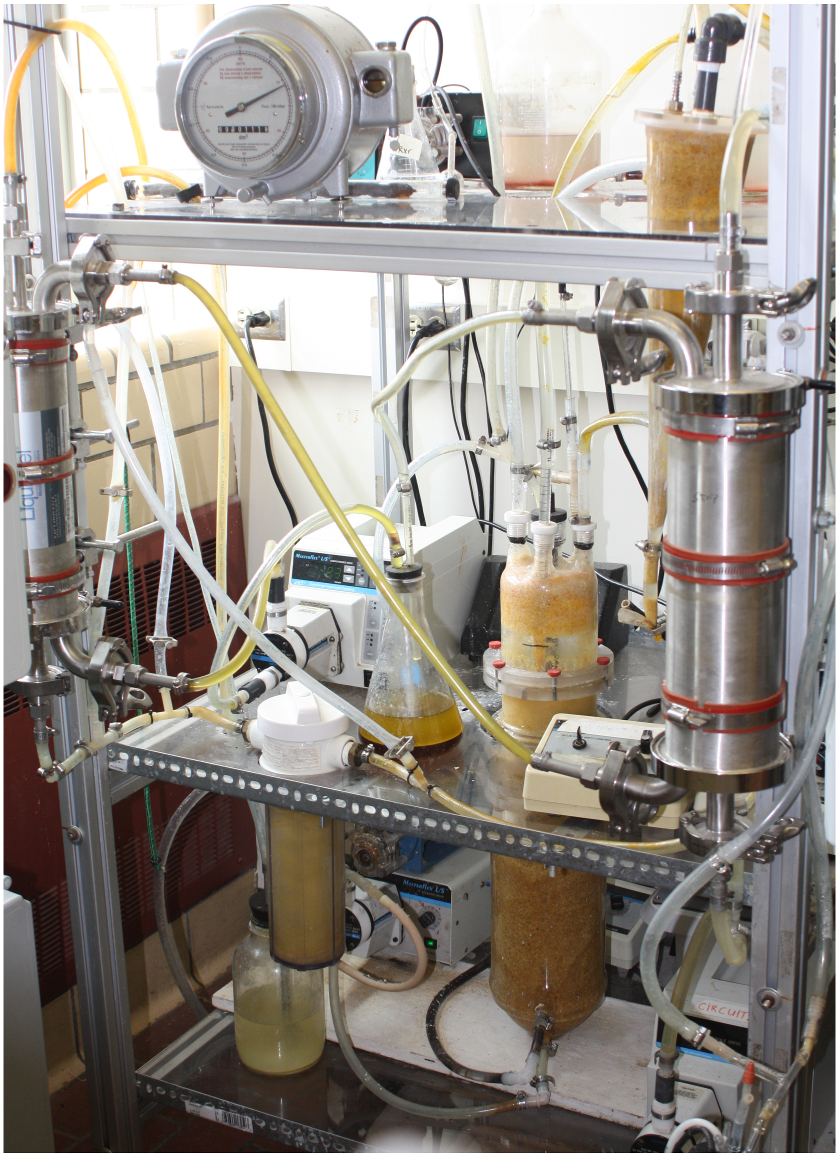

**Figure S2.** Picture of the bioreactor and membrane-based liquid-liquid extraction (pertraction).

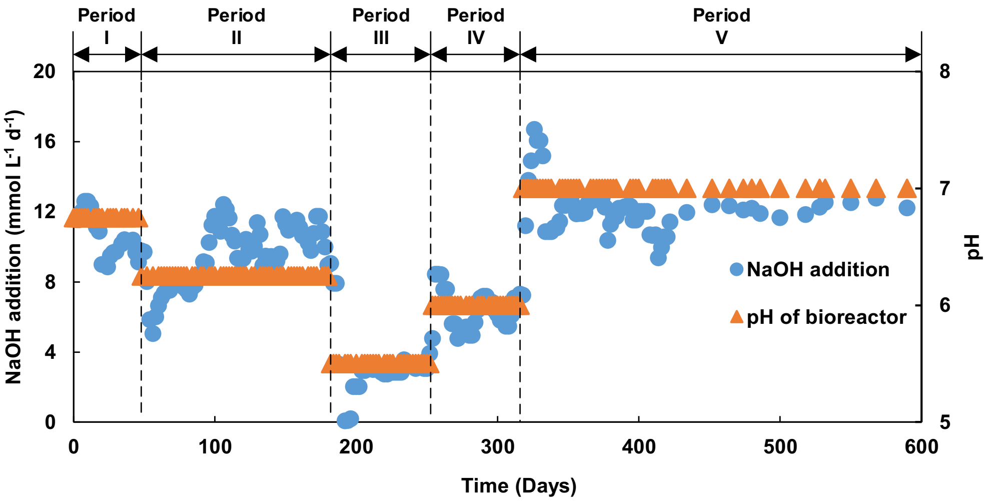

**Figure S3.** Time course of pH monitoring and NaOH addition during the operation of the *n*-caprylate-producing reactor.

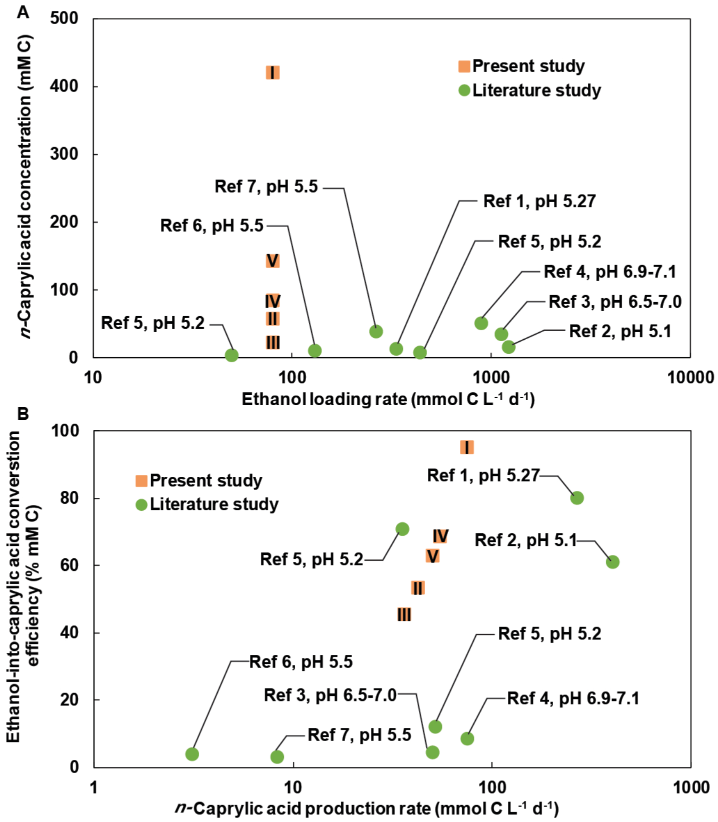

**Figure S4**. *n*-Caprylic acid production rate from the bioreactor with ethanol as an electron donor from published studies (green circles) and this study (orange squares). The published studies and this study: (A) *n*-caprylic acid concentration in the fermentation broth with ethanol loading rate; and (B) the ethanol-into-caprylic acid conversion efficiency and the *n*-caprylic acid production rate are listed from Ref 1,[1] Ref 2,[2] Ref 3,[3] Ref 4,[4] Ref 5,[5] Ref 6,[6] and Ref 7.[7] Operating periods of I to V in the present study are labeled in orange squares. Our study achieved a maximum concentration of *n*-caprylic acid in the broth and a maximum ethanol-into-caprylic acid conversion efficiency of 419.69 mM C and 94.95%, respectively, compared to published studies.

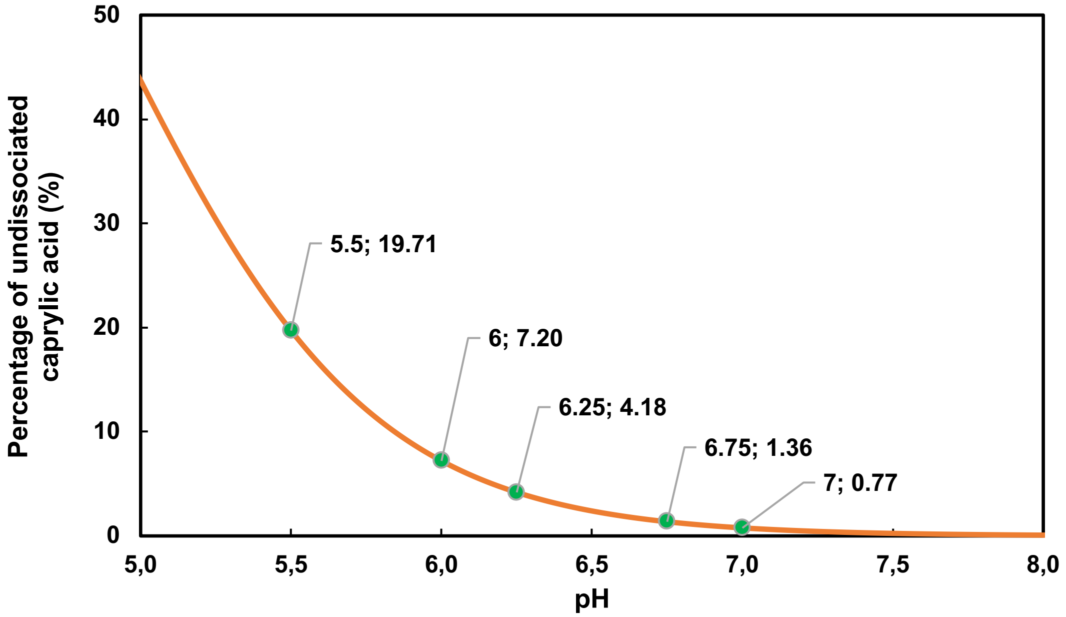

**Figure S5.** Relationship between undissociated *n*-caprylic acid in total *n*-caprylate at different pH values in aqueous solution.

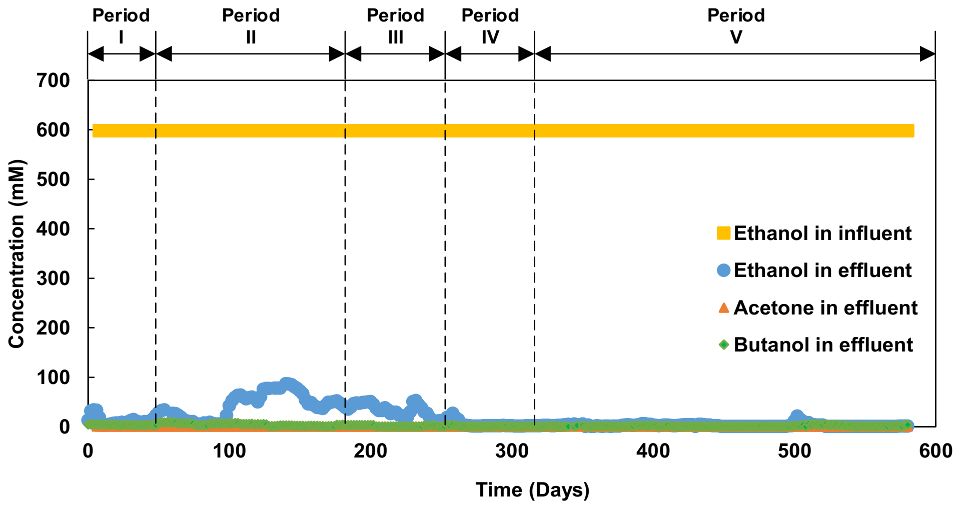

**Figure S6.** Time course of the concentration of ethanol, acetone, and butanol during the operation of the *n*-caprylate-producing reactor.

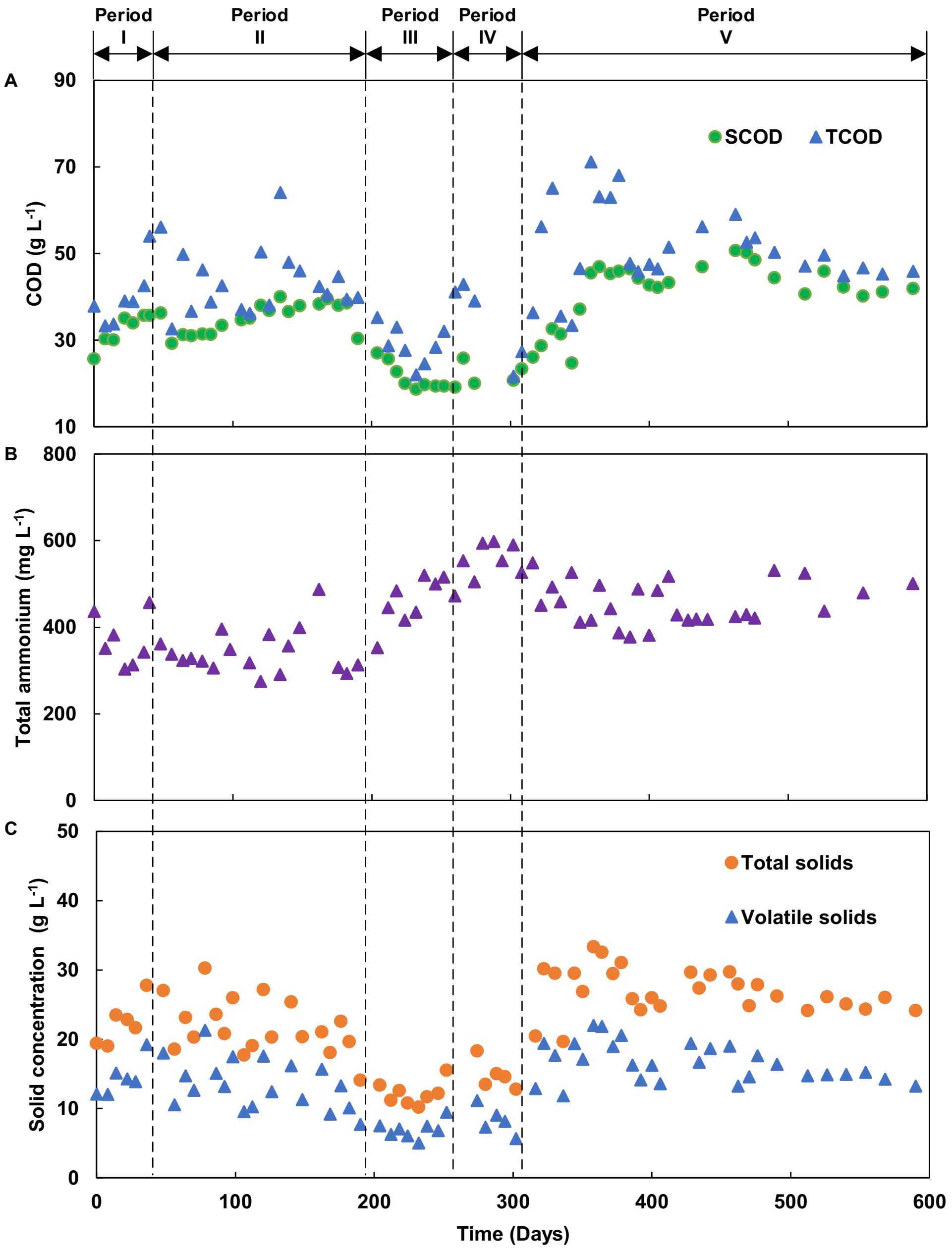

**Figure S7.** Time course of COD, ammonium, and solids data throughout the operating period. (A) SCOD and TCOD; (B) Total ammonium; (C) Total solids and volatile solids.

### Tables

**Table S1.** The composition of corn beer feedstock. Feedstocks were evaluated by analyzing six samples, and the error represents the standard deviation. Corn beer collected from Western New York Energy was used for all periods of the bioconversion system.

| Corn beer | pH | Ethanol ($g\ L^{-1}$) | Ethanol (mM C) | Ethanol ($g\ COD\ L^{-1}$) | Total COD ($g\ COD\ L^{-1}$) | Soluble COD ($g\ COD\ L^{-1}$) | Total Solids ($g\ L^{-1}$) | Volatile Solids ($g\ L^{-1}$) | Inert Solids ($g\ L^{-1}$) |
|---|---|---|---|---|---|---|---|---|---|
| Average | 4.60 | 122.21 | 5305.4 | 254.66 | 461.33 | 350.40 | 121.833 | 109.69 | 12.15 |
| STDEV | 0.0075 | 1.77 | 76.84 | 3.67 | 18.15 | 11.70 | 1.46 | 1.01 | 2.26 |

* COD conversions are: 0.096 g COD $mmol^{-1}$ for ethanol.

**Table S2.** Experimental approaches and conditions for batch methanogens inhibition.

| **Groups** | **pH** | **Undissociated *n*-caprylic acid (mM)** | **Groups** | **pH** | **Undissociated *n*-caprylic acid (mM)** | **Groups** | **pH** | **Undissociated *n*-caprylic acid (mM)** |
|---|---|---|---|---|---|---|---|---|
| 1 | 7.0 | 0 | 6 | 6.0 | 0 | 11 | 5.5 | 0 |
| 2 | 7.0 | 0.154 | 7 | 6.0 | 0.154 | 12 | 5.5 | 0.154 |
| 3 | 7.0 | 0.308 | 8 | 6.0 | 0.308 | 13 | 5.5 | 0.308 |
| 4 | 7.0 | 0.462 | 9 | 6.0 | 0.462 | 14 | 5.5 | 0.462 |
| 5 | 7.0 | 0.770 | 10 | 6.0 | 0.770 | 15 | 5.5 | 0.770 |
